# Structure-based development of new cyclic compounds targeting PSD-95 PDZ3 domain

**DOI:** 10.1101/2023.08.10.552828

**Authors:** Mandar T. Naik, Nandita Naik, Tony Hu, Szu-Huan Wang, John Marshall

**Affiliations:** Department of Molecular Biology, Cell Biology and Biochemistry, Brown University, Providence, Rhode Island, 02912, United States of America; Department of Molecular Pharmacology, Physiology and Biotechnology, Brown University, Providence, Rhode Island, 02912, United States of America

**Keywords:** Structure-based drug design, NMR, PSD-95, Depression, Angelman syndrome

## Abstract

Aberrant BDNF signaling has been proposed to contribute to the pathophysiology of depression and other neurological disorders such as Angelman syndrome. We have previously shown that targeting the TrkB / PSD-95 nexus by peptidomimetic inhibitors is a promising approach for therapeutic intervention. Here we used structure-based knowledge to develop a new peptidomimetic compound series that fuses SynGAP-derived peptides to our prototype compound CN2097. These compounds target the PSD-95 PDZ3 domain and adjoining αC helix to achieve bivalent binding that results in up to 7-fold stronger affinity compared to CN2097. These compounds were designed to improve CN2097 specificity for the PDZ3 domain and limited SAR studies have been performed to improve their resistance to proteolysis.

---

Major depressive disorder (MDD) affects one out of five individuals in their lifetime and is a serious disability impacting millions of people across all ethnicities and socioeconomic backgrounds [1]. Conventional antidepressant drugs that increase monoaminergic signaling have a delayed onset of action. The NMDA receptor antagonist esketamine produces rapid antidepressant effects in conventional treatment-resistant patients [2], although its effects tend to be short-lived and psychotomimetic in nature with little to no benefit for cognitive restructuring [3]. Thus, there is an unmet need for the development of safe and rapid-acting antidepressants that result in long-lasting therapeutic benefits. Multiple theories have been formulated to explain the development of MDD [4]. According to the neurotrophin hypothesis, depression is associated with reduced expression or function of brain-derived neurotrophic factor (BDNF). Abnormal BDNF signaling has been proposed to underlie the pathophysiology of MDD as well as bipolar disorder [5]. The hippocampus is one of several limbic structures implicated in the pathophysiology of depression, where BDNF and its high-affinity receptor tropomyosin receptor kinase B (TrkB) play a critical role in synaptic plasticity [6]. Many antidepressant drugs directly bind TrkB facilitating its synaptic localization and activation by BDNF [7]. Postsynaptic density protein-95 (PSD-95) is a synaptic scaffolding protein required for BDNF-induced signaling and is involved in a range of neurological disorders broadly classified under the depressive, autism-spectrum, and neurodevelopmental classes [8].

PDZ domains play important roles in cell signaling from the formation and stability of protein complexes to relaying extracellular stimuli detected on transmembrane receptors to the intracellular pathway. PSD-95 contains three PDZ domains (named PDZ1, PDZ2, and PDZ3) that bind to the PDZ-binding motif (PBM) present on the C-terminal of the partner protein with a consensus sequence of -x-S/T-x-Φ, where Φ is a hydrophobic and x is nonspecific residue [9]. These domains show selective preference towards the partners they recruit. For example, PBMs from Kv1.4 and NR2B bind strongly to the PDZ1 and PDZ2 domains and not to the PDZ3 domain, while the PBM from protein CRIPT binds specifically to the PDZ3 domain [10]. The selective anchoring of a partner to a particular PDZ domain can be explained by the differences in the PBM sequence [11] and the binding specificity can be altered by post-translational modifications [12].

The crystal structure of the PDZ3 complex with CRIPT PBM [13] prompted Spaller and colleagues to perform rigorous structure-based optimization of the CRIPT sequence. They showed that a β-Alanine stabilized 19-member ring structure between the two non-specific ’x’ positions of the CRIPT PBM results in the strong PDZ3 binding cyclic peptide [14]. However subtle changes in the peptide composition can alter PDZ binding affinity, and the cyclic peptide, unlike its linear counterpart, also bound the PDZ1 domain [15]. Further studies on the CRIPT peptide testing eight different bridging strategies [16] and nine substitutions on the terminal valine residue [17], resulted in a modest improvement in the PDZ3 binding affinity partly due to the lack of a 3D complex structure to guide the peptides designed for these experiments. The composition of the CRIPT linear peptide [18] was also rigorously analyzed for stronger binding sequences by extensive screening of 278 organic-modified peptide library [19] and by molecular dynamics simulations [20]. The improvement from linear peptide optimization has resulted in sub-micromolar range binders, however, the cyclized CRIPT peptide has several advantages over its linear counterparts. It not only has comparable affinity but also is less susceptible to proteolysis and hence is more stable in the blood plasma.

We have developed a series of peptidomimetic compounds that achieve efficient delivery of the CRIPT cyclic peptides to the brain by different blood-brain barrier (BBB) crossing strategies and identified a compound named CN2097 based on the disulfide-linked chimera of the CRIPT cyclic peptide with a poly-arginine BBB crossing moiety. CN2097 decreases PSD-95 interaction with the protein Arc and promotes BDNF-induced TrkB/PSD-95 complex formation [21]. The exact molecular mechanism of the CN2097 mode of action is not fully understood but it has a strong potential to reverse the hyperexcitability of wide dynamic range neurons in the dorsal horn of neuropathic rats [22] as well as facilitate the induction of long-term potentiation in Angelman syndrome (AS) mice [21]. CN2097 has neuroprotective properties [23], and it can reduce the post-traumatic synthesis of proinflammatory mediators and attenuate long-term potentiation impairment [24]. Our work on the widely used chronic mild stress and chronic social defeat stress depression models highlights the efficacy of CN2097 as an anti-depression treatment [25]. We have recently demonstrated that CN2097 normalizes autophagy and can mitigate cognitive and motor dysfunction in AS mice [26]. Similarly, other groups have shown the utility of peptide-based therapeutics such as Tat-NR2B9c [27] and AVLX-144 [28] that target the PSD-95 PDZ1 and PDZ2 domains as promising neuroprotectants in stroke [29]. This body of data suggests that peptidomimetic compounds targeting PDZ domains have a strong potential to be developed as effective therapeutic agents for the treatment of a range of neurological disorders. Here, we describe our progress in the structure-based development of CRIPT cyclic peptide towards a new bivalent inhibitor series named Syn3.

## Materials and methods

### Expression and purification of the PDZ3 domain

The genes expressing PSD-95 PDZ1 domain (residues M61-P153), PDZ2 (residues E156-Q266), and PDZ3 domain (residues D300-E401), were subcloned in-frame with a non-cleavable C-terminal His-tag of the pET28 vector. The PDZ3cα protein (residues R309-S422) was prepared using a plasmid received from Prof. Mingjie Zang that expressed an N-terminally His-tagged protein followed by PreScission protease cutting site [30]. All recombinant proteins were expressed in *E. coli* BL21 Star (DE3) cells grown at 37^°^C in LB or M9 medium supplemented with 50 μg/ml kanamycin. Uniformly ^15^N- or ^13^C,^15^N-labeled samples were prepared by growing bacterial cultures in M9 minimal medium using ^15^NH4Cl and ^13^C-glucose as the sole nitrogen and carbon sources. The expression of different proteins was achieved by the addition of 0.5 mM IPTG during the mid-log phase of culture growth and the cells were harvested after 4 hours of IPTG induction. The bacterial pellet was disrupted using a sonicator and the proteins were purified using a nickel-NTA affinity column. The PDZ3cα samples were further digested with PreScission protease to release the His-tag. The final purification was performed by size-exclusion chromatography with a HiLoad 16/60 Superdex 75 column using pH 7.5 FPLC buffer containing 50 mM potassium phosphate, 100 mM NaCl, and 0.5 mM EDTA. The purity of recombinant protein samples was checked by SDS-PAGE and protein concentrations were calculated using 280 nM UV absorbance.

### Peptide synthesis

All peptides and peptidomimetic compounds used in this study were custom synthesized by WuXi AppTec and purified to >95% using reverse phase HPLC. The compound identity was confirmed by mass and NMR spectroscopy analysis. Compounds were provided as lyophilized powder in vacuum-sealed vials and stored at -20^°^C. Fresh stock solutions were prepared by reconstituting weighed quantities of the peptide with the FPLC buffer and the peptide concentrations were measured using 280 nM UV absorbance.

### Nuclear Magnetic Resonance (NMR)

NMR data were acquired on Bruker 850, 600, or 500 MHz NMR spectrometer using 5 mm glass NMR tubes. The backbone resonance assignments of free PDZ3 and PDZ3cα were confirmed with the 3D-HNCACB experiment. Systematic titrations of various compounds were followed by a series of 2D-HSQC spectra. The NMR assignments of Syn3-bound PDZ3 complexes were derived from the analysis of these HSQC spectra that showed the movement of peaks due to the fast exchange on the NMR timescale and were further confirmed by 3D-HNCA.

### Surface Plasmon Resonance (SPR)

SPR data were acquired on the Biacore X100 instrument at 25°C using a constant 30 μl/min flow of running buffer containing 50 mM potassium phosphate (pH 7.5), 100 mM NaCl, 0.5 mM EDTA, and 0.05% P20 surfactant. The PDZ3cα was immobilized on the ligand channel of the CM5 chip using the EDC/NHS protocol followed by ethanolamine treatment. Each SPR experiment consisted of 3 baseline buffer runs followed by 8 sample runs at different analyte concentrations. 30 μl analyte peptide was injected for 60 seconds using an autosampler to observe the association phase followed by the continued flow of running buffer for 90 seconds to observe the dissociation phase. The SPR chip was regenerated after each dissociation step using 30 μl of 10 mM glycine, pH 2.5, followed by 2 minutes of equilibration with running buffer. The SPR sensorgrams were analyzed using a 1:1 binding model using Biacore evaluation software.

## Results and Discussion

### CN2097 binds strongly to the PDZ3 domain

Structure-based design has been at the forefront of the development of peptidomimetic compounds that target tandem PDZ domains of PSD-95. The C-terminal sequence of protein CRIPT is selective in binding to the PDZ3 domain, and a cyclic peptide derived from that sequence is at the heart of compound CN2097 developed by us [21]. The cyclic peptide has a stronger affinity for the PDZ3 domain than the linear peptide, but it also binds to the PDZ1 domain that shares only 44% sequence identity. We studied the binding of CN2097 to all three PDZ domain samples by NMR. As reported earlier [15], the compound binds the PDZ1 domain (Fig. 1A and 1C) but at the identical equimolar ratios, the chemical shift perturbation was much smaller compared to the PDZ3 domain binding. CN2097 did not show detectable binding to the PDZ2 domain but elicited very strong perturbation in the backbone amide resonances of the PDZ3 domain (Fig. 1B and 1D). Even though the cyclic peptide was developed to bind the PDZ3 domain and shows a relatively higher affinity and stronger NMR chemical shift perturbation for the PDZ3 domain, an NMR solution structure of the cyclic peptide in complex with the PDZ1 domain has previously been solved (Fig. 1E) [15]. The binding interface seen in this structure is illustrated in Figure 2A. The overall similarity in the perturbation profiles of PDZ1 and PDZ3 domains indicates CN2097 also binds the PDZ3 domain between the α-helix and β2 strand with the two PBM defining residues, Thr4 and Val6, being critical for this interaction. The peptide cyclization constrains these residues in an optimal binding pose and is important for the reduced proteolysis of CN2097 in blood plasma. Based on our prior work on the other PDZ binding compounds, it was inferred that a peptidomimetic with selective binding to the PSD-95 PDZ3 domain will be beneficial for neurological pathways that shape cognition. However, the PDZ1:CRIPT cyclic peptide complex structure has limited utility towards the development of a PDZ3 selective agent as the compound binding interfaces are not fully conserved. The PDZ3 domain has a much longer β2:β3 connecting loop and it lacks a non-binding, but adjacent α-helix seen in the PDZ1 domain that is disrupted by the Pro346 residue of the PDZ3 domain. Moreover, the PDZ3 domain has an allosteric regulator αC helix after its C-terminus that is not present after the PDZ1 domain (Fig. 1E). The αC helix is important for a hidden allosteric regulation of PBM binding [31]. The CRIPT linear PBM binds a longer PDZ3cα fragment carrying the αC helix with 21-fold higher affinity than the isolated PDZ3 domain [31].

**Figure 1.**
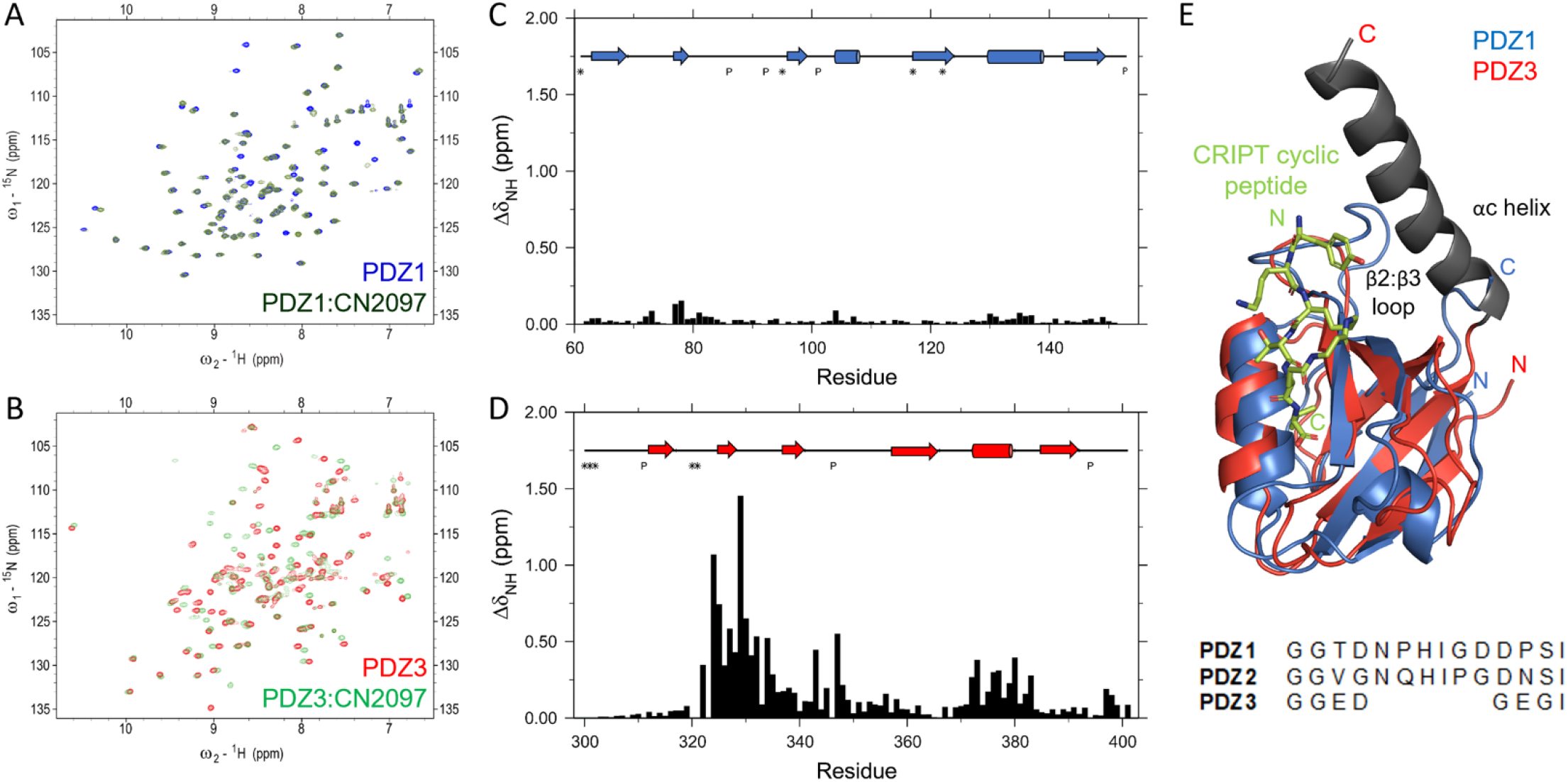
**A**. Overlay of 2D-^15^N-HSQC spectra of PDZ1 domain (blue) and 1:1 complex of PDZ1 and CN2097 (dark green), and **B**. PDZ3 domain (red) and 1:1 complex of PDZ3 and CN2097 (light green). This data was acquired at 20°C and pH 7.5. **C and D** are the corresponding chemical shift perturbations in the backbone amide resonances of PDZ1 and PDZ3 domains. The secondary structure of each domain and unobservable residue positions are marked at the top. **E**. Overlay of PDZ3 domain on PDZ1:CN2097 complex structure (1RGR). The sequence alignment of the β2:β3 loop region is shown below the ribbon representation created in Pymol. The αC helix (gray) was not present in the samples used for NMR studies on CN2097.

**Figure 2.**
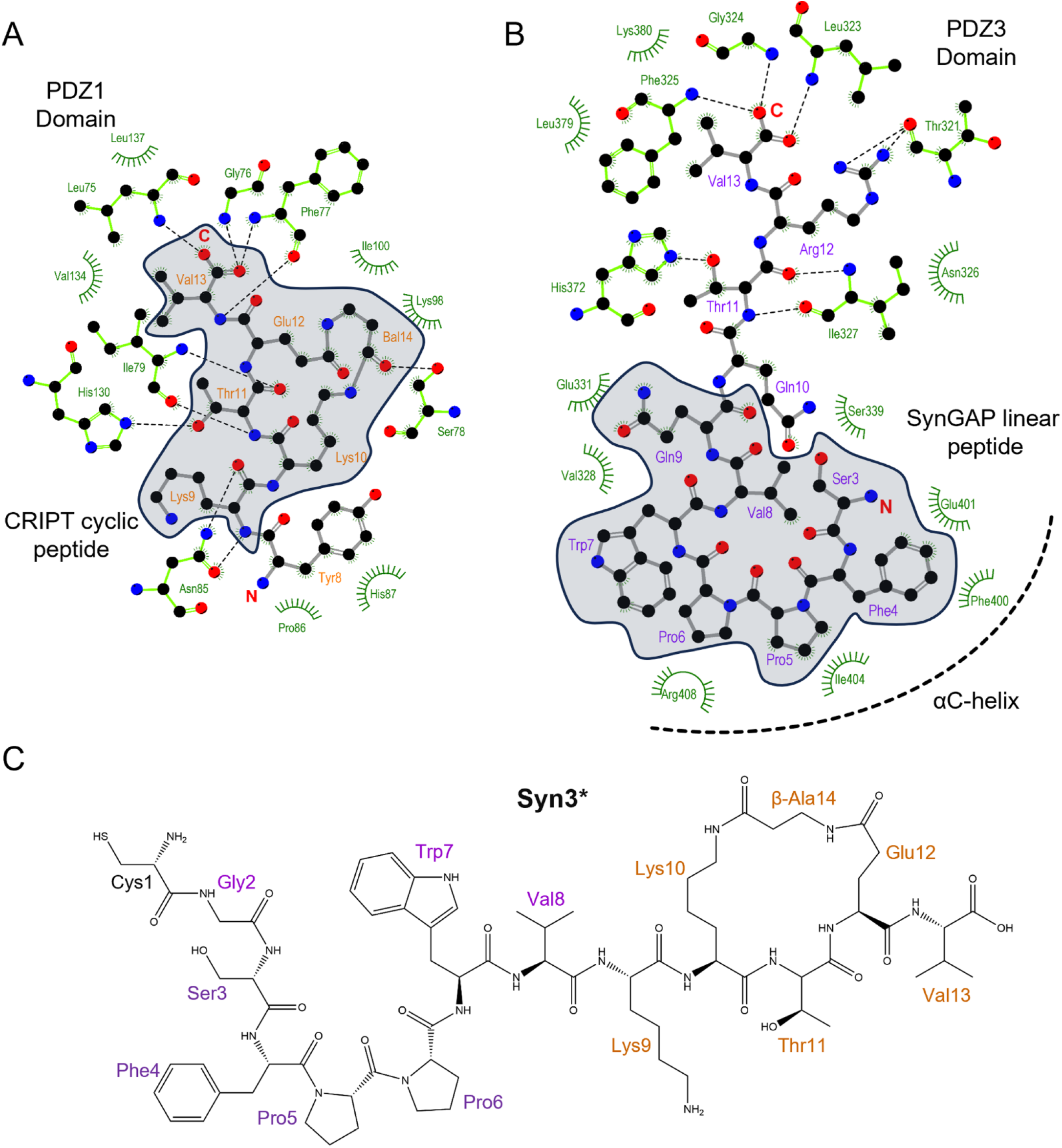
Binding interactions seen in **A**. the solution structure of the complex between PSD-95 PDZ1 domain and CRIPT cyclic peptide (PDB: 1RGR), and **B**. crystal structure of SynGAP linear peptide bound to PDZ3cα domain (PDB: 5JXB). Both peptides are shown with a ball and stick model while interacting PSD-95 residues are shown in green color with only electrostatically interacting residues being visible and hydrophobic interactions marked with curves. The peptide residues shown within the gray area were considered for the development of the Syn3 series. The peptide residues are renumbered differently than the original structure coordinates and the missing sidechain of Arg10 was reconstructed using UCSF Chimera [34]. The figure was created using the software Ligplot+ [35]. **C**. Structure of the active component peptide Syn3*. Syn3 is a disulfide-linked chimera of BBB crossing poly-arginine moiety and Syn3* (Table 1).

### Structure-based development of new candidates

Recently, we performed a detailed analysis of all PDZ3-bound structures available in the PDB database, none of which are cyclized peptides. We found that the PBM from SynGAP utilizes the αC helix present only after the PDZ3 domain, which was elegantly demonstrated by Zeng et al [30]. The interaction of SynGAP PBM with the longer PDZ3cα domain is shown in Figure 2B. The linear PBM peptides from CRIPT and SynGAP interact in a similar fashion with the folded PDZ3 domain due to the inherent recognition of the PBM, however, there are key differences. Unlike the CRIPT PBM, the SynGAP PBM does not distinguish the isolated PDZ3 domain from PDZ1 and PDZ2 domains but has a 15-fold higher affinity to the longer PDZ3cα fragment [10, 30]. The binding of SynGAP CC-PBM region to the PSD-95 PSG domain induces phase separation of the binary complex in liquid droplets reminiscent of the postsynaptic density and this phase separation behavior can be disrupted by a 15 residue peptide from SynGAP PBM but not by the CRIPT PBM [30]. These differences can be attributed to the N-terminus of SynGAP peptide directly binding to the allosteric regulator αC helix. In particular, hydrophobic interactions are seen in the crystal structure between the PBM residues Phe4, Pro6, and Trp7, and the αC helix. Here we have numbered the SynGAP residues differently to be consistent with the residue numbering used for the compounds we developed. In addition, phosphorylation by CaMKII of the SynGAP residue Ser1283, equivalent to the Ser3 residue of the peptide under our study, reduces binding affinity for the PDZ domain by 10-fold [32]. After consideration of the available structural information, we designed a hybrid compound named Syn3Q that fuses 8 helix-interacting residues from the SynGAP PBM at the N-terminus of the 5 residues from the CRIPT-derived cyclic peptide. This compound carried a cysteine at N-terminal to allow a covalent disulfide bond connection with the poly-Arg BBB crossing moiety. We then refined this peptide by combining 7 residues from SynGAP and 6 residues from the CRIPT cyclic peptide to generate Syn3. Both these peptidomimetic compounds were tested in an AS mouse model, where they mitigated impaired BDNF signaling to facilitate learning (data to be reported elsewhere). Our SPR experiments showed that Syn3 has a dissociation constant (K_D_) of 41 nM which is a 5-fold improvement over CN2097 for binding to the PDZ3cα (Table 1). SPR results also showed that both Syn3 and Syn3Q have higher on-rates *(k*_*a*_) as well as off-rates *(k*_*d*_) than CN2097. We performed NMR experiments that confirmed Syn3, similar to CN2097, binds the PDZ3 domain between the α-helix and β2 strand (Figure 3A-D). The chemical shift perturbation induced by Syn3 binding on the PDZ3 domain is very similar to CN2097 (Figure 1D) indicating the CRIPT cyclic portion binds the groove between β2 strand and α-helix. The strongest perturbation was seen for three glycines, Gly324, Gly329, and Gly330 (Fig. 3B-D) that tentatively interact with the PBM residues Thr11 and Val13. More importantly, Syn3 also shows above-average perturbation of the αC helix residues Ile404 and His405 that tentatively interact with the SynGAP-derived residues incorporated in Syn3. This bivalent binding is significant as it not only improves the affinity of CRIPT-derived cyclic compounds but also makes Syn3 more specific for binding to the PDZ3 domain as the PDZ1 domain does not have a helical element after its C-terminus. More significantly, Syn3 binding may interfere with the PDZ3 domain allosteric mechanism [31, 33] as suggested by the presence of complicated slow dynamics revealed by significant linewidth broadening observed in the PDZ3cα:Syn3 complex spectrum (Fig. 3B). The NMR study on Syn3 was conducted at 15°C to observe weak peaks for the amides of Gly324, Gly329, and Gly330 (Figure 3B) but almost every peak, except the Thr321, had weaker intensity in the Syn3 complex spectrum compared to the free PDZ3cα spectrum. Interestingly, the peak for Thr321 which was barely visible in the free-form spectrum became sharp in the complex. The loss in signal intensity indicates linewidth broadening of the resonances which could be a result of conformational exchange. The linewidth broadening did not affect CN2097 complex spectrum to the same extent even though they were acquired at a higher 20°C temperature. A detailed NMR relaxation study is required to probe the presence of ms-μs range motions and their potential role in the inhibition of the PDZ3 allosteric mechanism.

**Table 1.** SPR measurements on the binding of various peptides to the immobilized PDZ3cα.

| Peptide | Sequence | $k_a$<br>($10^5 \cdot M^{-1} \cdot s^{-1}$) | $k_d$<br>($10^{-3} \cdot s^{-1}$) | $K_D$<br>(nm) |
| --- | --- | --- | --- | --- |
| CN2097 | RRRRRRRC=CKNYK[KTE(β-Ala)]V | 0.06 | 1.09 | 190.6 |
| Syn3Q | RRRRRRRC=CGSFPPWVQ[KTE(β-Ala)]V | 2.61 | 31.16 | 119.4 |
| Syn3 | RRRRRRRC=CGSFPPWVK[KTE(β-Ala)]V | 17.75 | 73.23 | 41.3 |
| Syn3* | CGSFPPWVK(KTE(β-Ala))V | 4.39 | 28.70 | 65.3 |
| D-Syn3 | (d-Arg)x7.(d-Cys)=(dCys)GSF(d-Pro)PWVQ[KTE(β-Ala)]V | 4.58 | 12.54 | 27.4 |
| <u>SynGAP control peptides</u> |  |  |  |  |
| SYNGAP | RRRRRRRCCGSFPPWVQQTRV | 0.22 | 0.65 | 29.3 |
| αC binder | GSFPPWVQ | 0.01 | 5.92 | 11111.0 |
| <u>L- amino acid substitutions</u> |  |  |  |  |
| V8L | RRRRRRGSFPPWLQQTRV | 0.65 | 3.81 | 58.4 |
| V8I | RRRRRRGSFPPWIQQTRV | 0.29 | 2.94 | 100.8 |
| F4H | RRRRRRGSHPPWVQQTRV | 2.16 | 26.09 | 120.8 |
| <u>Modified L- amino acid substitutions</u> |  |  |  |  |
| F4-NH2F | RRRRRRGS(ζ-AminoPhe)PPWVQQTRV | 1.53 | 10.40 | 67.8 |
| W7-OHW | RRRRRRGSFPP(5-hydroxy-Trp)VQQTRV | 0.24 | 10.64 | 445.5 |
| <u>D- amino acid substitutions</u> |  |  |  |  |
| D-Ser3 | RRRRRRG(d-Ser)FPPWVQQTRV | 6.33 | 4.93 | 7.8 |
| D-Pro4 | RRRRRRGSF(d-Pro)PWVQQTRV | 18.81 | 11.56 | 6.1 |
| D-Pro5 | RRRRRRGSFP(d-Pro)WVQQTRV | 18.28 | 44.35 | 24.3 |
| D-Val8 | RRRRRRGSFPPW(d-Val)QQTRV | 21.90 | 45.86 | 20.9 |
| D-Gln9 | RRRRRRGSFPPWV(d-Gln)QTRV | 3.17 | 18.17 | 57.3 |

**Figure 3.**
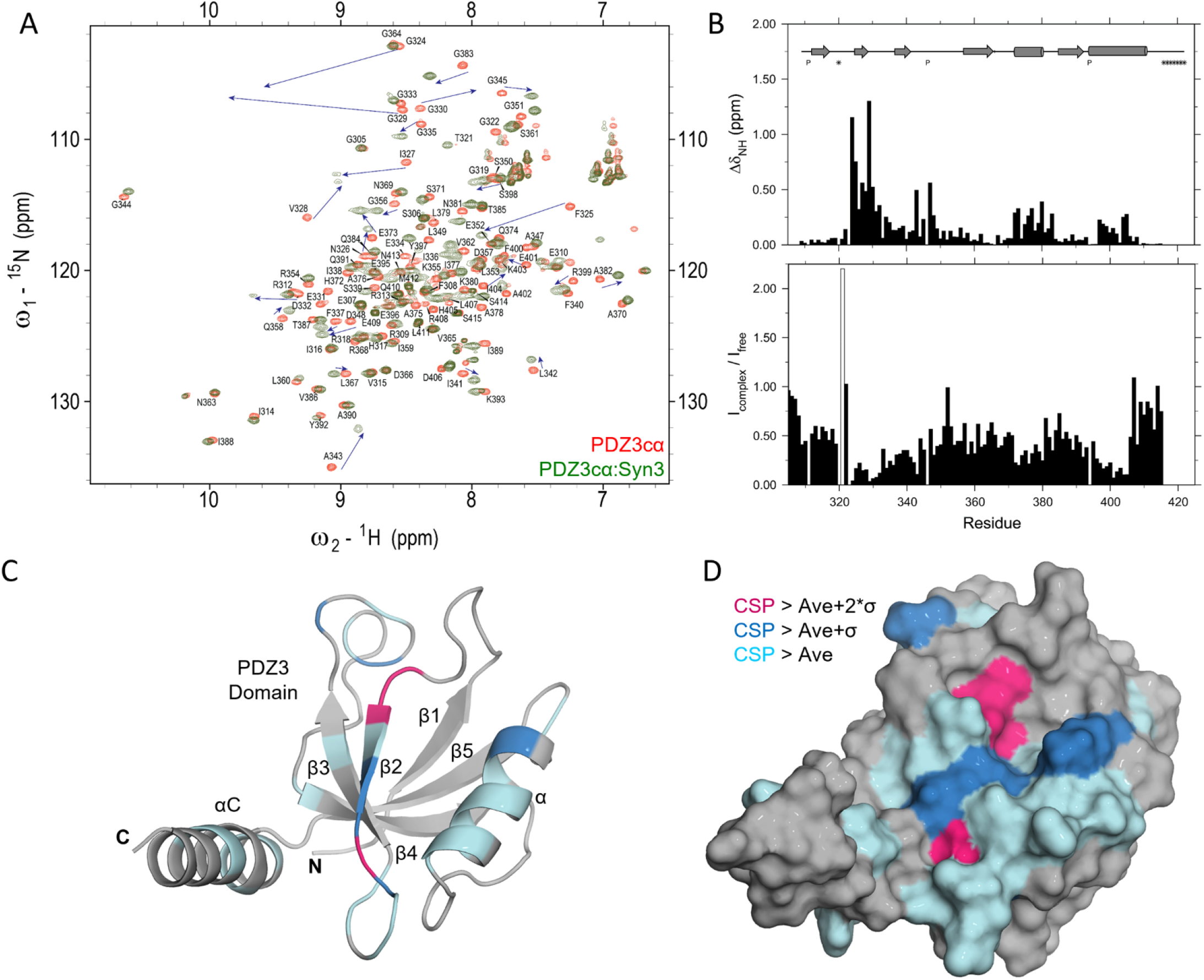
**A**. Overlay of 2D-^15^N-HSQC spectra acquired at 15°C and pH 7.5 on PDZ3cα (red) and its 1:1 complex with Syn3 (olive green). The peaks are annotated with the assignments for free PDZ3cα and the movement of strongly perturbed peaks is shown with arrows. **B**. The average chemical shift perturbation (above) and the ratio of peak intensities (below) for individual PDZ3 backbone amides are shown as bar plots. The peak intensity ratio for T321 is shown with an open bar. **C**. Color-coded ribbon and **D**. surface representation of PDZ3cα structure (PDB: 5JXB). This figure was prepared using Pymol [36].

### Structure-activity relationship optimization of Syn3

We then tested the original linear SynGAP-derived peptide, without the cyclic component but with a poly-Arg tag, for PDZ3cα binding. Surprisingly, under identical assay conditions, this SYNGAP peptide bound with a 2-fold higher affinity than Syn3 and had a slower off-rate, however, it did not have the same beneficial effects as Syn3 in our animal studies. A smaller fragment of this peptide made up of only αC helix interacting residues was also tested and found to have a weaker dissociation constant of 11 μM. These results indicate that the αC helix provides an avenue for synergistic binding enhancement for the PBM and there is further scope for improvement in the affinity of the Syn3 compound series. As it was more cost-effective to study linear peptides than cyclic compounds, we then tested ten knowledge-based modifications to the SYNGAP peptide sequence that included changes with 3 regular L-amino acid mutations, 2 substitutions with non-natural amino acids, and 5 substitutions with D-amino acids. In addition, F4E W7A and V8E, substitutions had been previously reported in the literature [30]. We identified that changing Pro4 to a D-analogue improves binding affinity by 2-fold. Since D-amino acids are less susceptible to proteolysis, they are desirable in peptidomimetic drugs. We then synthesized a new analogue of Syn3 that carried Pro5 substituted with a d-Proline residue as well as substituting the entire arginine tag and cysteines with D-amino acids. This compound was named D-Syn3 and has a comparable binding affinity to the linear SYNGAP peptide for the PDZ3cα domain. The Syn3 and D-Sy3 analogues have been tested in animal models and the results of those experiments will be presented elsewhere.

In summary, through a rigorous analysis of structures of other PDZ-binding proteins, we have developed a new compound series that consists of a cyclic peptide identical to CN2097 but is augmented with amino acids derived from SynGAP that bind to the extended αC helix of PDZ3. Additional structural and biophysical characterization is required to fully understand the atomic details of compound interaction with the PDZ3 domain and elucidate the effects on PDZ3 allosteric mechanism. These studies are also critical for the further development of the CRIPT peptidomimetic series.

## Abbreviations

AS: Angelman syndrome;
BBB: blood-brain barrier;
BDNF: brain-derived neurotrophic factor
CSP: chemical shift perturbation;
MDD: major depressive disorder;
PBM: PDZ-binding motif;
PSD-95: postsynaptic density protein-95;
TrkB: tropomyosin receptor kinase B.

## Author contributions

MN and JM designed the study; MN, NN, TH, and SW performed the experiments and analyzed the data; MN and JM wrote the manuscript.

## Acknowledgments

This research is based in part on data obtained at the Brown University Structural Biology Core Facility, which is supported by the Division of Biology and Medicine, at Brown University. This work was supported by generous funding from the National Institutes of Health R01NS094440, R21MH104252, and STTR R41MH118747, the Harrington Discovery Institute, and the Foundation for Angelman Syndrome Therapeutics.

